# Cell-type specific inference from bulk RNA-sequencing data by integrating single cell reference profiles via EPIC-unmix

**DOI:** 10.1101/2024.05.23.595514

**Authors:** Chenwei Tang, Quan Sun, Xinyue Zeng, Xiaoyu Yang, Fei Liu, Jinying Zhao, Yin Shen, Bixiang Liu, Jia Wen, Yun Li

## Abstract

Cell type specific (CTS) analysis is essential to reveal biological insights obscured in bulk tissue data. However, single-cell (sc) or single-nuclei (sn) resolution data are still cost-prohibitive for large-scale samples. Thus, computational methods to perform deconvolution from bulk tissue data are highly valuable. We here present EPIC-unmix, a novel two-step empirical Bayesian method integrating reference sc/sn RNA-seq data and bulk RNA-seq data from target samples to enhance the accuracy of CTS inference. We demonstrate through comprehensive simulations across three tissues that EPIC-unmix achieved 4.6% - 109.8% higher accuracy compared to alternative methods. By applying EPIC-unmix to human bulk brain RNA-seq data from the ROSMAP and MSBB cohorts, we identified multiple genes differentially expressed between Alzheimer’s disease (AD) cases versus controls in a CTS manner, including 57.4% novel genes not identified using similar sample size sc/snRNA-seq data, indicating the power of our *in-silico* approach. Among the 6-69% overlapping, 83%-100% are in consistent direction with those from sc/snRNA-seq data, supporting the reliability of our findings. EPIC-unmix inferred CTS expression profiles similarly empowers CTS eQTL analysis. Among the novel eQTLs, we highlight a microglia eQTL for AD risk gene *AP3B2*, obscured in bulk and missed by sc/snRNA-seq based eQTL analysis. The variant resides in a microglia-specific cCRE, forming chromatin loop with *AP3B2* promoter region in microglia. Taken together, we believe EPIC-unmix will be a valuable tool to enable more powerful CTS analysis.

## Introduction

Gene expression, commonly measured from bulk tissue samples by RNA-sequencing (RNA-seq) technology, is important for quantitative trait loci study and downstream genetics research ^1–3^. Though such bulk tissue transcriptomic profiles can reflect the etiology of some diseases, such as Alzheimer’s disease (AD), to a certain degree, they cannot capture functional heterogeneity across cell types due to the mixture of multiple cell types. Therefore, cell-type-specific (CTS) analysis, which can reveal biological insights obscured in the bulk tissue data, has become increasingly popular ^4–6^. One gold-standard to obtain CTS gene expression is from single-cell (sc) or single-nuclei (sn) RNA-seq data. However, such single-cell technology is still cost-prohibitive for population samples and may not even be feasible. For example, currently the largest snRNA-seq data for AD has 427 individuals ^7^; but besides this, the second largest study included only 24 cases and 24 controls ^6^. In addition, such sc/snRNA-seq data also tend to be noisier than bulk RNA-seq data due to technical difficulties, limiting their potential to be directly used in genetic analysis. As an alternative, researchers have been relying on computational methods for CTS analysis.

With the increasing availability of bulk RNA-seq data generated from tissue samples, there is a unique opportunity for CTS inference by integrating bulk RNA-seq data and the burgeoning CTS profiles derived from a group of relatively homogenous cells in the sc/snRNA-seq data, which is known as “deconvolution”. There have been many computational deconvolution methods developed in the past few years, and we group them into two categories. Most of the methods, especially in early years, estimate only the cell-type fractions of bulk tissues for each study sample rather than the sample-level CTS expression profiles, including CIBERSORT ^8^, MuSiC ^9^, and Bisque ^10^. We classify them as “traditional” deconvolution methods (**Figure S1**). The second category more ambitiously aims to estimate CTS expression profiles separately for each sample. The output for these methods are k sample-by-gene matrices, where k is the number of cell types, starting from one sample-by-gene matrix at the bulk level. We name them as “aggressive” methods (**Figure S1**). Currently, there are few methods falling into this category, only including CIBERSORTx ^11^, TCA ^12^ and bMIND ^13^. CIBERSORTx ^11^ is a machine learning method that uses non-negative least squares to infer sample-level CTS expression with the aim of grouping samples. TCA ^12^, originally designed for methylation data, can also be adapted for RNA-seq based expression data. It is a frequentist method that does not employ any reference in the CTS inference step. Both CIBERSORTx and TCA missed the opportunity to borrow information from sc/snRNA-seq data. In contrast, a Bayesian method bMIND ^13^ takes sc/snRNA-seq reference data to build its prior, but this method is susceptible to the differences between reference and target samples, leading to unstable results for different datasets.

To address these challenges, we developed EPIC-unmix, EmPirical bayes cell type specifIC unmixing of bulk expression profiles, a two-step Bayesian method integrating sc/sn RNA-seq reference data and bulk RNA-seq data from targeted samples to enhance the accuracy of CTS expression inference. We performed comprehensive simulation studies using data representing different human and mouse tissues to evaluate the performance of EPIC-unmix, where the results showed that EPIC-unmix achieved the best performance for almost every scenario. Encouraged by the simulation results, we then performed real data analysis to deconvolute bulk RNA-seq data from Religious Orders Study/Memory and Aging Project (ROSMAP) ^14^ and Mount Sinai Brain Bank (MSBB) ^15^, carrying out a gene selection strategy validated from the simulations. We conducted meta-analyses based on results derived from the two datasets, and identified some differentially expressed genes between AD cases and controls, followed by CTS eQTLs and cCREs. We believe that our EPIC-unmix offers a new paradigm to explore the CTS mechanism of complex diseases.

## Result

### Overview of EPIC-unmix

EPIC-unmix is a two-step empirical Bayesian method that takes sc/snRNA-seq data as reference to perform CTS deconvolution while accounting for the difference between reference and target samples (**Figure 1**). In the first step, EPIC-unmix employs sc/snRNA-seq as a reference to infer CTS expression, using the same Bayesian framework adopted by the published bMIND method ^13^. The major improvement of EPIC-unmix compared to bMIND lies in the second step where EPIC-unmix adds another layer of Bayesian inference based on the prior derived from the CTS expression inferred for the target samples in the first step (**Methods**). The second layer of Bayesian inference renders the EPIC-unmix model data adaptive, as it can adjust for the difference between reference and target datasets. Such a data adaptive step makes EPIC-unmix conceptually more stable than bMIND.

**Figure 1.**
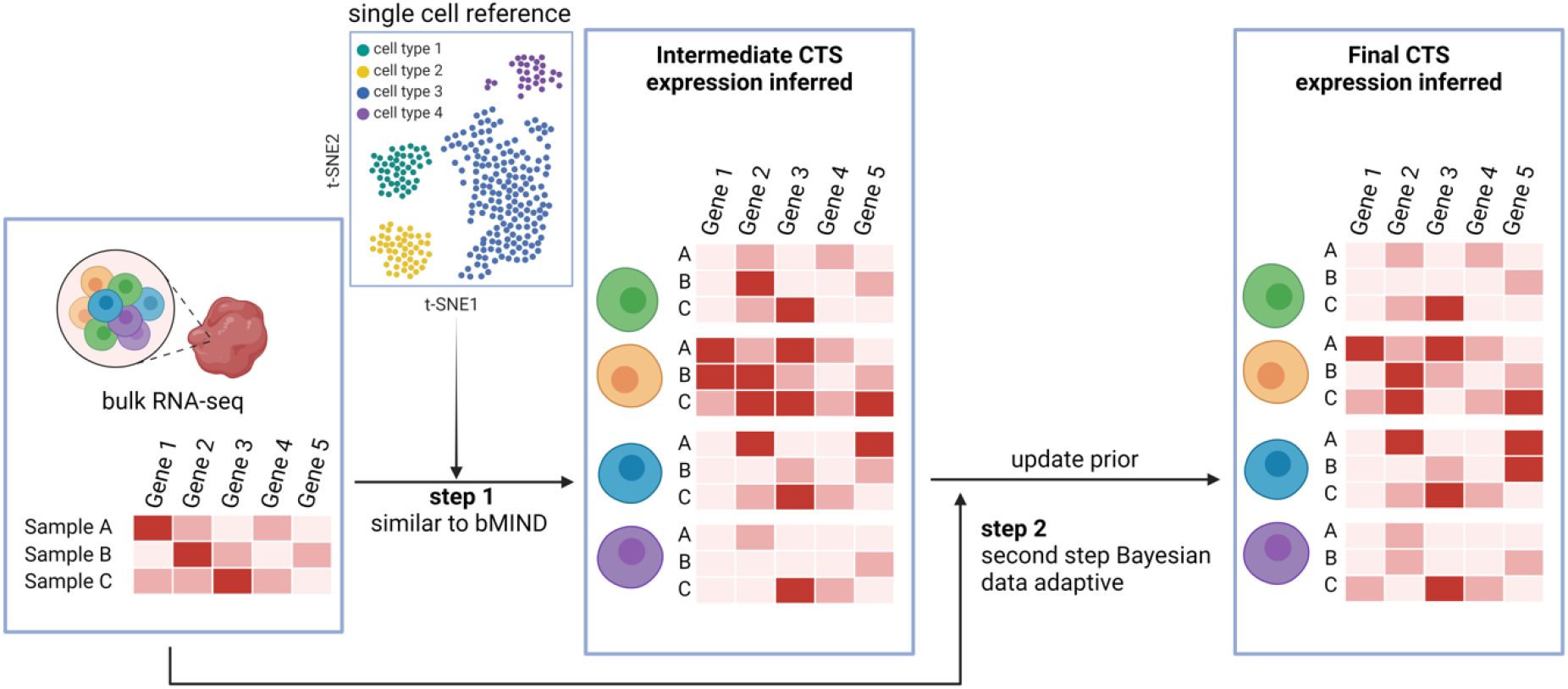
Overview of EPIC-unmix. EPIC-unmix is a two-step empirical Bayesian method. In the first step, it takes sc/snRNA-seq data as reference to perform Bayesian inference for target bulk RNA-seq data, similar to bMIND. After obtaining the CTS profiles, EPIC-unmix continues to the second step, where it updated the prior information from the derived CTS profiles in the first step and performed a second round of Bayesian inference. By design, the updated prior contains information from both the original sc/snRNA-seq reference and also the target, which is thus data adaptive and can account for potential discrepancies between reference and target in terms of donor characteristics and other biological or technical confounders.

### EPIC-unmix achieves better performance in ROSMAP human brain dataset

We first performed real data based simulations leveraging snRNA-seq data from ROSMAP ^16^ (n = 48) to evaluate the performance of EPIC-unmix in comparison with two previous published methods, TCA ^12^ and bMIND ^13^. In brief, we removed one sample that failed quality control (QC), and then randomly selected 16 samples as targets and generated pseudo-bulk, with the remaining 31 (10 case, 21 control) samples as references (**Methods**). Since not all genes could be deconvoluted with high accuracy, we developed a gene selection strategy based on cell-type marker genes and agreement among sc/snRNA-seq datasets and CTS bulk RNA-seq data (**Methods**). Specifically, it combines multiple sources including external brain snRNA-seq data ^17^, cell type specific marker genes from the literature ^18,19^, as well as marker genes inferred from an internal ROSMAP snRNA-seq dataset ^16^ and the bulk RNA-seq data ^20^. The final gene list includes 1003, 1916, 764, and 548 genes for microglia, excitatory neurons, astrocytes, and oligodendrocytes respectively (**Table S1**).

Comparing deconvolution accuracy for the selected gene set versus unselected genes in simulations, we observed that selected genes demonstrated better performance than unselected genes in each of the four cell major types (**Figure 2A**). For example, using EPIC-unmix, selected genes showed 45.2% and 56.9% higher mean and median Pearson Correlation Coefficient (PCC) compared to unselected genes across all the cell types (Wilcox signed-rank test p-value < 1e-4). Such advantages were similar when comparing different reference panels (psychENCODE scRNA-seq ^21^ and ROSMAP snRNA-seq data^16^) and deconvolution methods (**Figure S2**), indicating the robustness of our gene selection strategy. Additionally, EPIC-unmix achieved 4.6% - 66.1% and 3.2% - 25.3% higher median PCC compared to TCA (Wilcox signed-rank test p-value = 1.64e-55) and bMIND (Wilcox signed-rank test p-value = 5.86e-4), and the superiority of EPIC-unmix was again robust to the choice of reference panels and across multiple cell types (**Figure 2B**).

**Figure 2.**
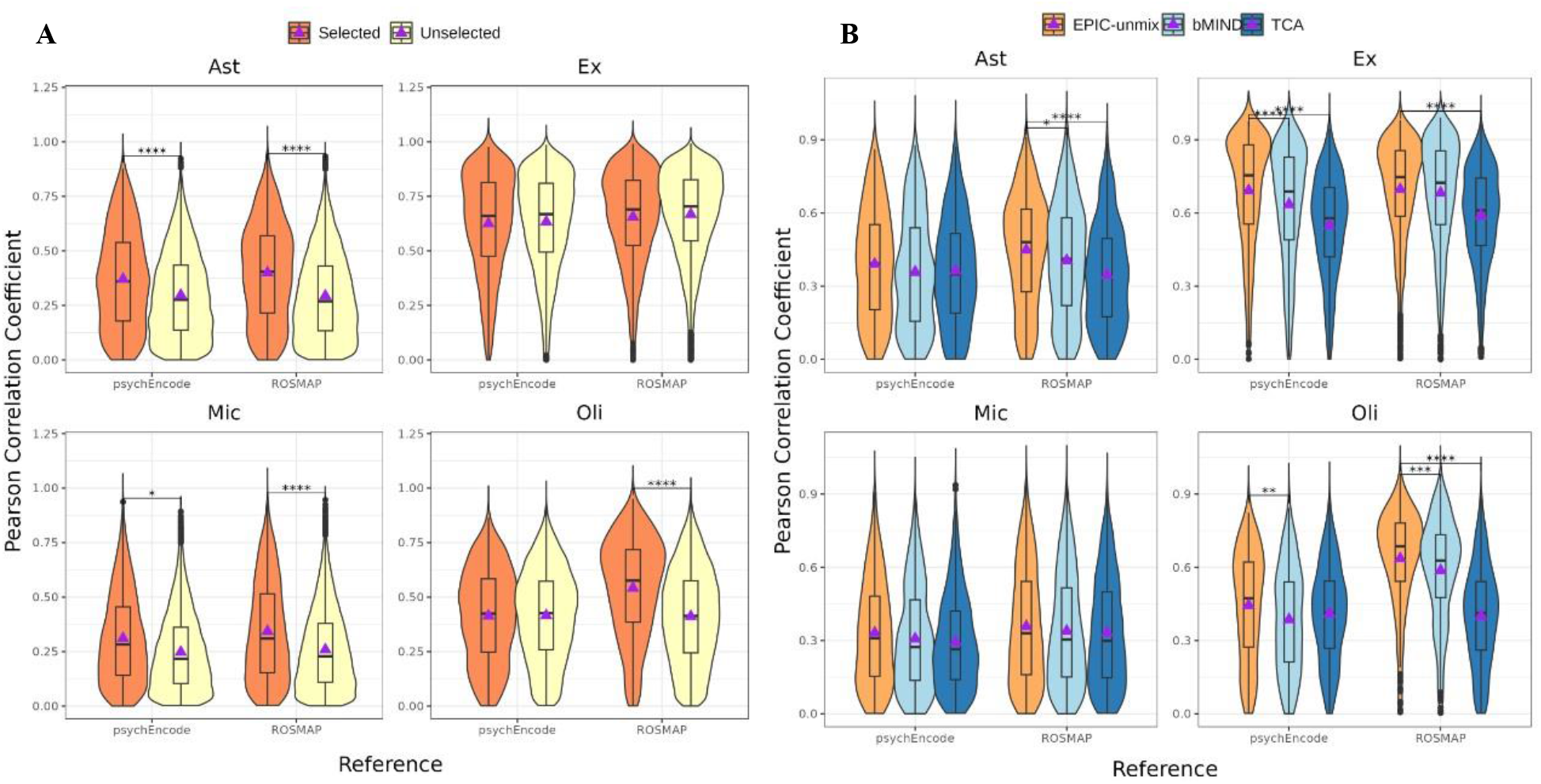
Performance comparison in ROSMAP-based simulations. **A**. Comparing performance for selected and unselected genes, with two different reference panels (psychENCODE, ROSMAP) along the X-axis. EPIC-unmix was applied for deconvolution. **B**. Comparing different deconvolution methods, similarly for two different reference panels (psychENCODE, ROSMAP) along the X-axis. Y-axis denotes the absolute Pearson correlation between inferred CTS expression and truth.

### EPIC-unmix shows stable and robust performance across multiple datasets

Encouraged by the promising results in the ROSMAP-based simulated brain data, we further evaluated the robustness of EPIC-unmix using more comprehensive simulations with a wide source of sc/snRNA-seq data from both mouse brain and human blood tissues, similarly compared to bMIND and TCA using multiple reference panels.

We first leveraged the scRNA-seq data including 116 cell types from primary motor cortex from Yao et al. ^22^ to generate pseudo bulks for half of the donors (n = 112) randomly selected. We applied EPIC-unmix, bMIND, and TCA to the pseudo bulks for deconvolution, using either the rest of single cell data from Yao et al. or an external mouse neocortex scRNA-seq dataset (Tasic ^23^) as reference for EPIC-unmix and bMIND. Similarly, we adopted a gene selection strategy based on cell-type marker genes (**Methods**) and observed better inference for selected genes in all cell types (**Figure 3A**). Encouragingly, we found that EPIC-unmix again showed the best performance among the three methods for all cell types (**Figure 3B, Table S2**). For example, in L5 PT, the mean PCC of EPIC-unmix was 15.7% and 109.8% higher than that of bMIND and TCA, respectively (Wilcox signed-rank test p-values < 1e-4 for both). In addition, we found similar advantage of our EPIC-unmix when performing deconvolution with an external scRNA-seq dataset from Tasic et al. ^23^ (**Figure 3B, Table S2**), suggesting the robustness of EPIC-unmix in terms of reference panels used.

**Figure 3.**
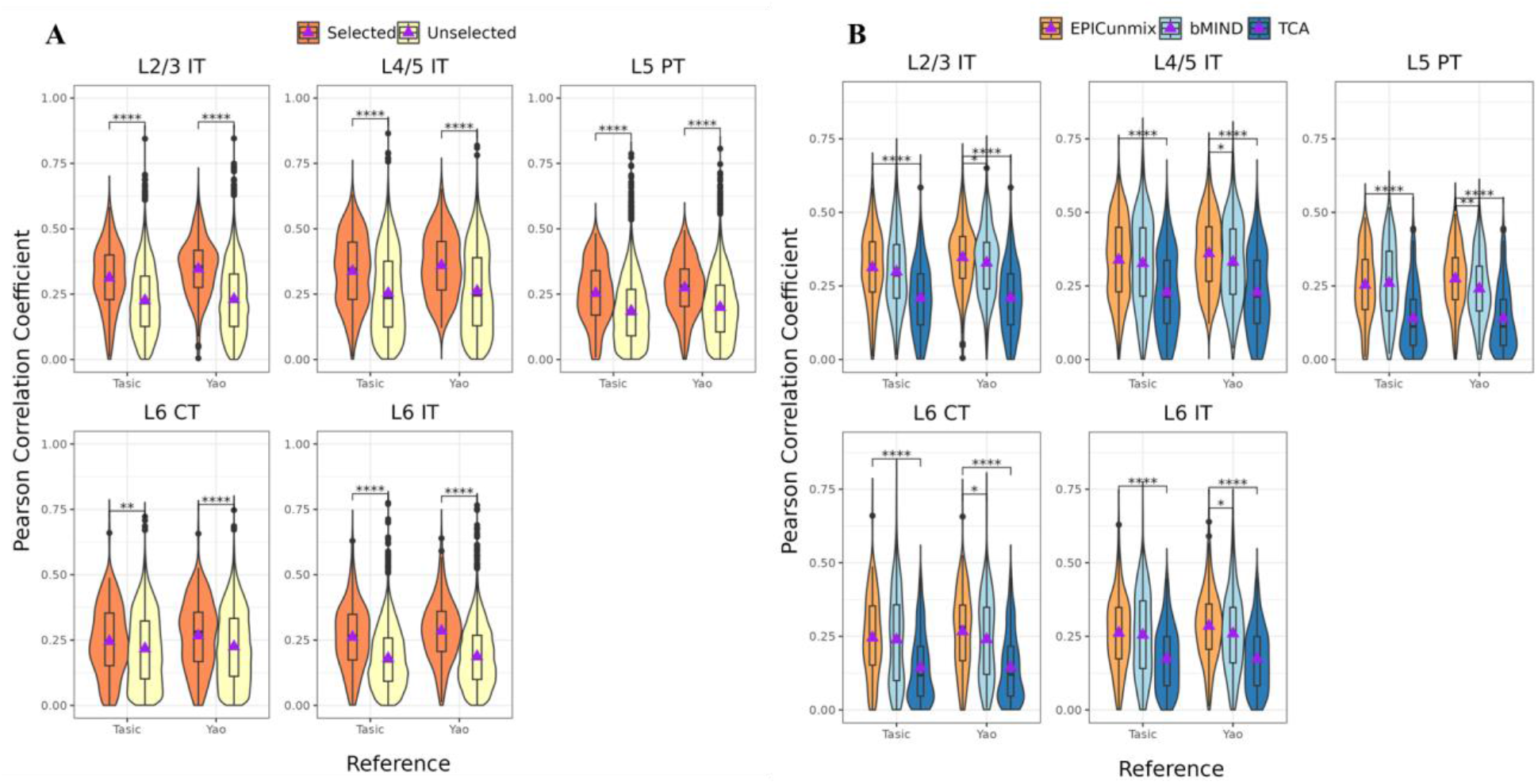
Performance comparison in mouse brain scRNA-seq-based simulations. **A**. For pseudo bulks generated from Yao et al 2021, we compared the correlation between true and EPIC-unmix estimated CTS gene expression with two reference panels (Tasic and Yao) of selected and unselected genes over five cell types. Y-axis denotes the absolute Pearson correlation between inferred CTS expression and truth. **B**. Performance of EPIC-unmix, bMIND, and TCA for selected genes. Similarly Y-axis indicates the absolute PCC, X-axis indicating reference panels. P-value significances were obtained from one tailed Wilcoxon signed-rank test of whether EPIC-unmix’s PCC had higher means.

Observing the superior performance of EPIC-unmix across multiple brain datasets for both human and mice species, we next proceeded to a different tissue. Specifically, we obtained scRNA-seq data of human PBMCs from the AIDA study ^24^ and similarly generated pseudo bulks from half of the donors (n = 117) randomly selected. We then performed deconvolution using as reference either the rest single cell data from AIDA or an external human blood scRNA-seq dataset (OneK1K ^5^). We found that EPIC-unmix achieved higher PCC than bMIND in all cell types, and higher than TCA for all except memory B cells (**Figure 4, Table S3**). As an example, the mean PCC of EPIC-unmix in natural killer cells were 105.1% and 54.9 % higher than that of bMIND and TCA, respectively (Wilcox signed-rank test p-values < 1e-4 for both). Note that when using the external OneK1K ^5^ as the reference, EPIC-unmix still showed the best overall performance among the three methods (**Figure 4, Table S3**), again indicating the robustness of EPIC-unmix under the realistic scenario of external reference panels. These simulation results show that EPIC-unmix is more stable and robust across different targeted bulk data and reference panels from different species and tissues than the existing two deconvolution methods.

**Figure 4.**
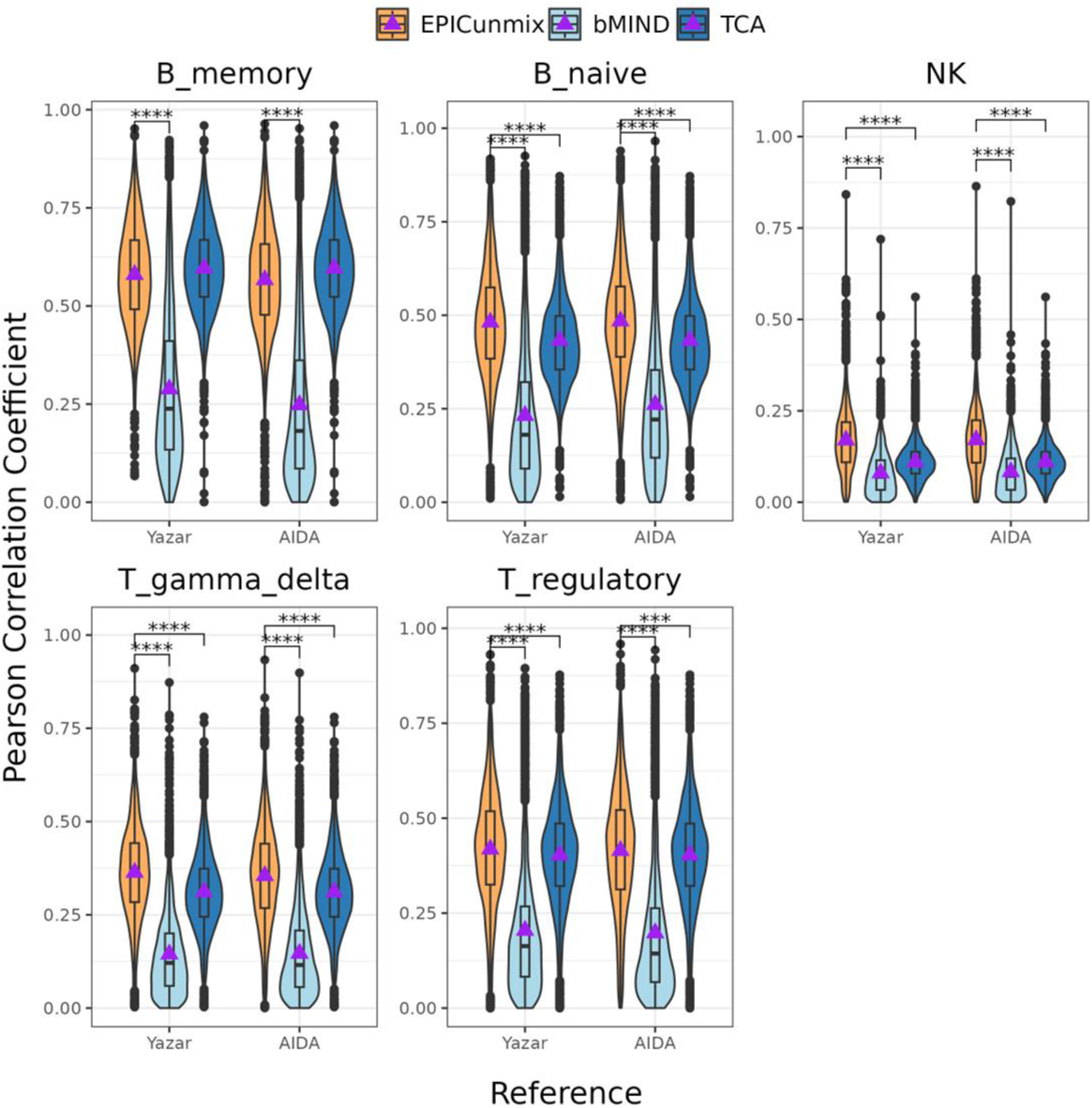
Performance comparison in human PBMC scRNA-seq-based simulations. For pseudo bulks generated from AIDA, correlation between true and estimated CTS gene expression over five cell types were computed to compare the performance of EPIC-unmix, bMIND, and TCA, with two different reference panels (along the X-axis) for EPIC-unmix and bMIND. Y-axis denotes the absolute PCC between inferred CTS expression and truth. The p-value significances were obtained from one tailed Wilcoxon signed-rank test of whether EPIC-unmix’s PCC had higher means.

### EPIC-unmix boosts power for downstream CTS differential expression and eQTL analysis

Encouraged by results from real data based simulations, we then applied EPIC-unmix to two human brain bulk RNA-seq data, ROSMAP ^14^ (n = 401) and MSBB ^15^ (n = 250), using snRNA-seq of 48 samples (no overlap with target samples) as reference for both datasets to perform deconvolution. After obtaining EPIC-unmix inferred CTS profiles, we performed differentially expressed gene (DEG) analysis between AD cases and controls in a CTS manner, adjusting for appropriate covariates (**Methods**), separately for the two datasets. We then performed a meta-analysis using METAL ^25^ combining the two sets of results.

We identified 14, 49, 30, and 28 DEGs in astrocytes (Ast), excitatory neurons (Ex), microglia (Mic) and oligodendrocytes (Oli) respectively, at a false discovery rate (FDR) of 5% (**Table S4**). Among microglia DEGs, we highlight *SLC6A12*, which was considered a hub gene with high expression in AD patients ^26^; and *ADAMTS2*, a potential therapeutic target for adult brain disorders including AD and schizophrenia from a previous study ^27^. We additionally performed a pathway analysis for microglia DEGs (**Methods**) and identified important biological pathways related to AD, including neurotransmitter uptake/reuptake ^28^ and tubulin-binding activity (**Table S5**). Comparing our results with DEGs derived from ROSMAP snRNA-seq data (n = 427) in a most recently published study ^7^, we identified 2-34 overlapping CTS DEGs, with 83-100% genes exhibiting consistent directions of effects (**Table S6, Method**).

We also carried EPIC-unmix inferred CTS gene expression profiles to perform CTS cis-eQTL analysis, similarly separately for ROSMAP and MSBB first, followed by meta-analysis (**Method**). Note that all the analyses were performed solely based on deconvoluted CTS gene expression profiles inferred by EPIC-unmix, without any snRNA-seq data involved. We discovered 63,043-110,292 CTS eQTLs (FDR < 5%) and 1,246-1,829 CTS eGenes (defined as genes with any eQTL at FDR < 5%) across the four cell types, representing 125,259 unique eGene-variant pairs and 2,210 unique genes. We compared our *in silico* CTS cis-eQTL results with a previous study based on scRNA-seq data from Bryois et al ^29^, n=192). We found that 12.9-26.4% of our significant CTS cis-eQTL gene-variant pairs were also nominally significant (p ≤ 0.05) in Bryois et al. For these shared gene-variant pairs, we observed that the z-scores of our eQTL results were significantly correlated with the effects sizes of those from Bryois et al in all four cell types, with 86.0-90.7% eQTLs showing consistent direction of effects and PCC ranging from 0.792-0.886 (**Figure 5A, Table S7**). For the remaining cis-eQTL exclusively identified in our results (but not nominally significant in Bryois et al), we compared the microglia specific ones with those from two microglia eQTL studies (Young et al (n = 93) ^30^ and Kosoy et al (n = 101)^31^), both using sorted primary microglia. We found that 23.4% and 14.7% of cis-eQTL gene-variant pairs exclusively identified by us were also significant in these two datasets respectively, while only 6.5% and 4.8% of exclusive microglia eQTLs from Bryois et al can be replicated by the two microglia eQTL studies.

**Figure 5.**
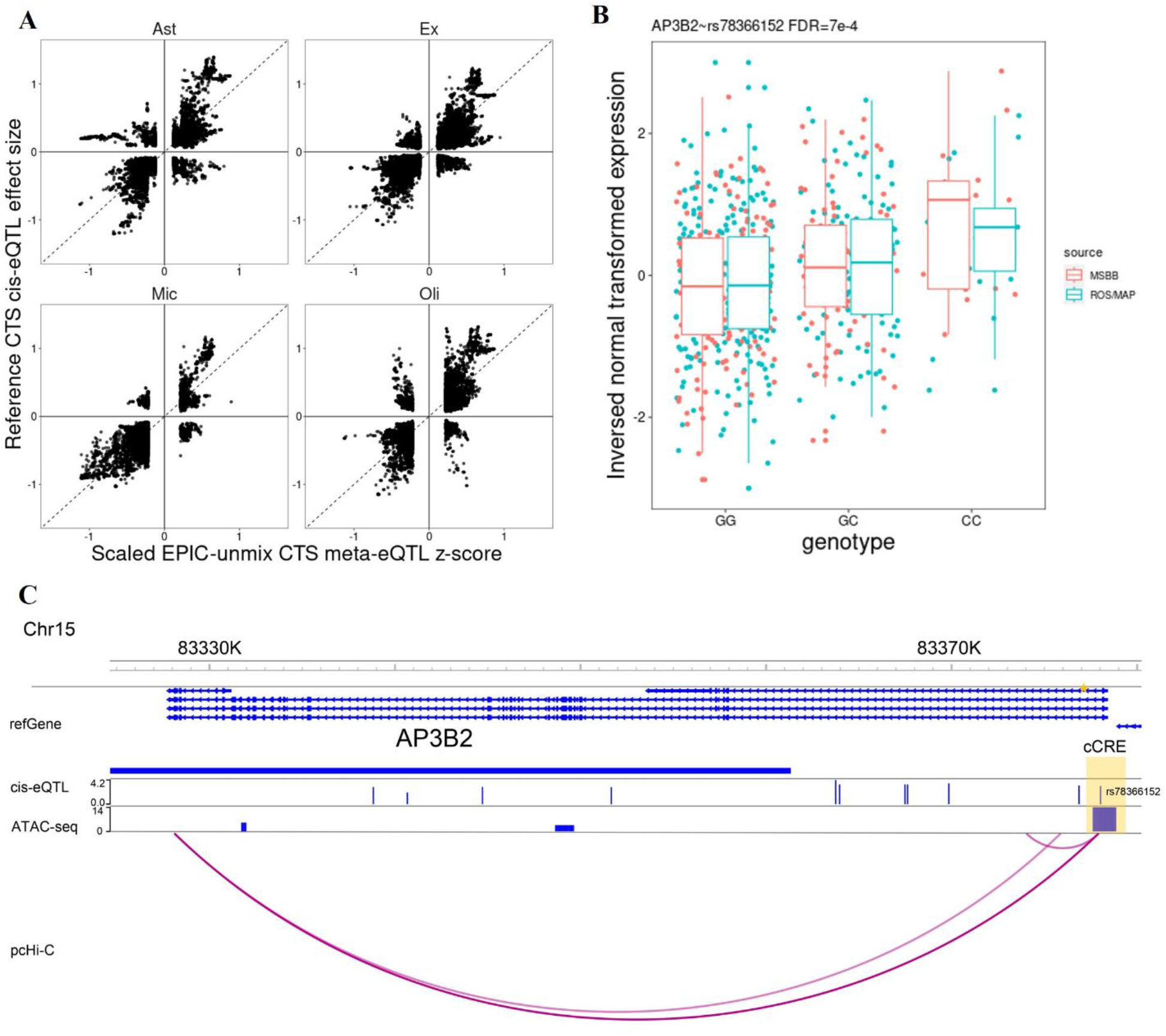
CTS cis-eQTL from EPIC-unmix deconvolution after meta-analysis. **A**. Effect size comparison of CTS cis-eQTL from EPIC-unmix deconvolution and snRNA-seq. X-axis is meta z-scores, scaled to range from -1 to 1, from EPIC-unmix; y-axis is effect sizes from Bryois et al. Only overlapping eQTLs with FDR < 5% in both studies are included. **B**. Genotype of cis-eQTL rs78366152 vs. EPIC-unmix inferred microglia gene expression. **C**. Annotated genome browser view of *AP3B2*. Tracks from top to bottom are: genome position with GRch37 build; gene location; *AP3B2*’*s* CTS cis-eQTL (height of signal = -10log(adjusted_eQTL_p)); microglia specific ATAC-seq narrow peaks (height of signal = score); microglia specific pcHi-C loops.

Notably, our results distinctly discovered a CTS cis-eQTL, rs78366152, for *AP3B2*, an AD risk gene interrogated in multiple mouse model studies ^32,33^ (**Figure 5B**). In our eQTL meta-analysis results, this gene-variant pair is significant in microglia, oligodendrocyte, and astrocyte, but not in excitatory neuron. Note that rs78366152 was not identified as an eQTL for *AP3B2* with ROSMAP bulk brain RNA-seq data ROSMAP^6^; nor was it reported as a microglia-specific cis-eQTL (**Table S7**). Moreover, rs78366152 resides in a microglia-specific ATAC-seq peak that forms a loop with *AP3B2* gene upstream region (**Figure 5C**) based on pcHi-C data ^20^, suggesting that rs78366152 might exert microglia-specific regulatory effect on *AP3B2* expression. Additionally, we highlight rs72825667, a microglia-specific eQTL of *MERTK*, a drug target and a signaling gene involved in various processes including macrophage clearance of apoptotic cells, platelet aggregation, cytoskeleton reorganization and engulfment ^34^, residing in its cCRE region (**Figure S3A**); and multiple microglia-specific eQTLs of *KAT8*, an AD associated gene ^35^ colocalizing with AD GWAS variants (**Figure S3B**) ^36^. These findings demonstrate EPIC-unmix can enable powerful downstream CTS analysis given only bulk RNA-seq data in study samples for biologically important discoveries in a cell-type-specific manner.

## Discussion

Cell type specific analysis is crucial to reveal biological insights and/or disease pathology, while before high-quality and sc/snRNA-seq data are available in large samples, computational methods are needed to facilitate such analyses. In this manuscript, we present EPIC-unmix, a reference based deconvolution method. It adopts a two-step Bayesian framework that incorporates information from sc/snRNA-seq reference in the first step while adaptively accounting for the differences between reference and targets in the second step. We performed comprehensive simulation studies based on multiple sc/snRNA-seq datasets from different tissues to compare EPIC-unmix with other state-of-the-art methods, demonstrating its outstanding performance and robustness to different target and reference datasets. To examine how EPIC-unmix could facilitate real data analysis, we applied EPIC-unmix to deconvolute bulk RNA-seq data using snRNA-seq from ROSMAP and MSBB, and carried forward the derived CTS profiles into downstream CTS DEG and CTS cis-eQTL analyses. Both analyses reported cell type relevant findings consistent with previously published results based on snRNA-seq data or sorted CTS data, while reporting novel findings. These results emphasize the ability of EPIC-unmix to provide computationally and biologically interpretable CTS expression profiles at individual sample level.

Our extensive simulations included different datasets from human brain, mouse brain and human blood tissues. While EPIC-unmix always achieved the best performance, the pattern between bMIND and TCA was not consistent (**Figure 2-4**). We observed bMIND achieved better performance over TCA for the two brain tissues, while TCA outperformed bMIND for human blood tissue. Such inconsistencies may be attributed to the underlying intrinsic differences between sc/sn references and target samples. For the two brain tissues, reference provided essential information and thus boosted CTS inference; while for the blood tissue, there may exist significant dissimilarities between donor characteristics or technical confounders, preventing bMIND to efficiently borrow information from the reference scRNA-seq. However, our EPIC-unmix, specifically thanks to its second step, could mitigate systematic differences between reference and target samples to improve CTS inference. Our results also highlight the robustness of EPIC-unmix to different reference panels.

In light of the fact that CTS expression cannot be perfectly inferred from bulk expression for every gene in every cell type, we designed gene selection strategies to prioritize genes that are most likely well inferred in each cell type. We retained the gene lists considering both gene expression patterns from the reference dataset and deconvolution quality from simulations, as well as cell type marker genes from previous studies. We observed that by restricting deconvolution to these most promising genes, EPIC-unmix as well as other deconvolution methods achieved better prediction accuracy (**Figures 2, S2**). Our practice demonstrates that, as an example, a proper gene selection strategy boosts the performance of CTS inference, likely benefiting from reduced low-quality input to the prediction model, which is analogous to feature selection frequently applied in various prediction and classification tasks in genetic studies ^37,38^. We note that we did not consider an uncertainty measurement of the deconvolution quality in this study due to its complexity, which warrants future explorations to leverage, for example, information from a pre-training stage that provides estimated quality for each gene and cell type combination.

In the CTS DEG analysis using our EPIC-unmix estimated CTS profiles and by meta-analyzing ROSMAP and MSBB datasets, we confirmed the reliability of our results by observing highly consistent results derived from sc/snRNA-seq in ROSMAP ^7^. Meanwhile, we recognize that our results are limited in power compared to snRNA-seq based studies if the sample size is similar or the same. We note that for most tissues, the number of samples with bulk tissue RNA-seq data is much larger than those with sc/snRNA-seq data, providing a big market for EPIC-unmix, before sc/snRNA-seq are routinely available for most study samples.

We also performed CTS eQTL analysis with the EPIC-unmix inferred CTS expression profiles, again using ROSMAP and MSBB data, and similarly observed reassuring results when comparing our cis-eQTL signals with those derived from snRNA-seq study from Bryois et al ^29^. Nevertheless, such comparisons (primarily focusing on p-values and effect sizes) to some extent oversimplifies the comparison due to underemphasis of context differences. For example, ROSMAP, MSBB and Bryosis et al sequenced similar but different brain regions, and the study populations also differ in many aspects, such as the compositions of AD case and controls, age, sex, and genetic ancestry etc. Additionally, we identified some novel eQTLs that were missed by previous studies. As an example, we highlighted one microglia eQTL of *AP3B2* located in a cCRE specific to the same cell type, strengthening the credibility of its regulatory effect on *AP3B2* gene expression.

We note that in our simulations and real data analyses, especially for ROSMAP and MSBB AD studies, we ignored AD status when performing deconvolutions, for both snRNA-seq reference and target samples with bulk tissue RNA-seq data only. We acknowledge the potential differences between cases and controls which may affect the inference. However, simple separate inferences for cases and controls may lead to more severe problems, including batch effects, smaller sample sizes for inference, and more importantly increased false discoveries due to manual separation in analysis of cases from controls. Future studies are warranted to explore better strategies to perform deconvolution in such case-control studies.

In summary, we developed EPIC-unmix, a two-step empirical Bayesian method integrating sc/sn RNA-seq reference and bulk RNA-seq data from targeted samples to enhance the accuracy of CTS gene expression inference. Through comprehensive simulations and real data analysis, we demonstrated the advantages of EPIC-unmix over alternative methods. We believe our method provides valuable insights to facilitate cell type specific analysis in larger sample sizes before sc/snRNA-seq data are massively available.

## Methods

### Overview of EPIC-unmix

Let *i* = 1, …, *N*; *j* = 1, …, *G*; and *k* = 1, …, *K* be the sample, gene and cell type indices respectively, where *N, G*, and *K* denote the total number of samples, genes and cell types. Further, let *X*_*ij*_ denote the log2-transformed bulk RNA-seq gene expression value of gene *j* for sample *i*, 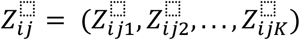 denote the *K* × 1 vector of cell-type-specific gene expression levels across the *K* cell types for gene *j* of sample *i*, and 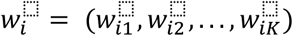 denote the *K* × 1 vector of cell type proportions for sample *i*. Similarly to TCA ^12^ and bMIND ^13^, we assume 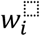 is known, which can be estimated using many existing methods, such as MuSiC ^9^, MuSiC2 ^39^, SCDC ^40^, Bisque ^10^ and CIBERSORT ^8^. Note that the purpose of our method is to derive individual-level CTS gene expression profiles rather than merely estimating cell type fractions.

Following the bMIND work ^13^, we similarly assume 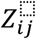 follows a multivariate normal distribution with mean 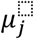 and covariance matrix *Σ*_*j*_, and the relationship between the observed bulk gene expression and the underlying CTS expression follows the model below:

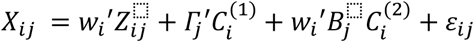

where 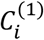 and 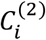 correspond to bulk- and CTS-specific covariates; 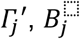 are the coefficients for 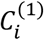 and 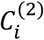, respectively; and 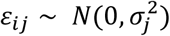 is the noise term.

In the first step of EPIC-unmix, we derive the priors of 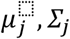 and 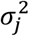 from sc/snRNA-seq reference data. Specifically, we assume 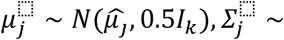 InvWishart 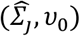 and 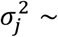 InvWishart (1, 0), where 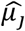 is the average CTS expression derived from reference (also at the log_2_ scale), 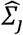 is the empirical covariance matrix also derived from reference, and *υ*_0_ controls the degree of belief of the reference, with higher value denoting higher confidence. We note that the inverse Wishart distribution is the conjugate prior for the multivariate normal distribution, and the prior for 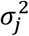 is a non-informative prior.

In the second step, we update the priors using the estimated CTS expression obtained from the first step, assuming the same distributional form, specifically 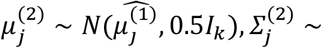 InvWishart 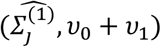 and 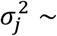 InvWishart (1, 0). The parameters in the update priors, namely 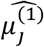 and 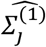, are calculated based on the estimated CTS expression for the target samples, with the rationale that such a prior is more compatible with the target samples and is less susceptible to the difference between reference and target samples. Specifically, 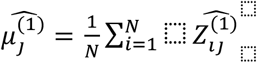, where 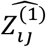 is the estimated CTS expression for gene *j* of the *i*th target sample after the first step, and 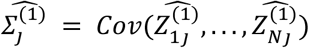 is the corresponding *K* × *K* sample covariance matrix for gene *j* based on the target samples. We note that the updated prior for 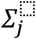 is slightly different from that in the first step. We add a non-zero term *υ*_1_ to the original parameter *υ*_0_ since the updated prior has now integrated information from the target samples, which is closer to the target samples than the original prior. Note that both *υ*_1_ and *υ*_0_ are pre-specified parameters.

Parameter estimation in both steps is achieved using Markov Chain Monte Carlo (MCMC) sampling implemented in R package “MCMCglmm” ^41^.

### Datasets

#### The ROSMAP human dorsolateral prefrontal cortex dataset

##### ROSMAP ^42^

Religious Orders Study (ROS) and the Rush Memory and Aging Project (MAP) (collectively, ROSMAP) data include bulk RNA-seq data and snRNA-seq data, both from the dorsolateral prefrontal cortex. The availability of both bulk and snRNA-seq data provides the opportunity to perform CTS deconvolution using EPIC-unmix. Bulk RNA-seq data includes 638 samples, while earlier snRNA-seq data includes only 48 individuals (24 cases and 24 controls) with a total of 70,634 cells across multiple cell types including but not limited to microglia, astrocytes, excitatory neurons, oligodendrocytes. Among the 638 samples, we retained only those with the physician’s overall cognitive diagnostic category (cogdx) being 4 or 5 (treated as AD cases) or 1 (treated as controls), leaving a final set of 441 samples ^42^ in this study

##### MSBB ^15^

Mount Sinai/JJ Peters VA Medical Center Brain Bank (MSBB) includes bulk RNA-seq data across 4 brain regions (the frontal pole, superior temporal gyrus, parahippocampal gyru and inferior frontal gyrus) and whole genome sequencing data (WGS) for 364 individuals. We used the bulk RNA-seq data from the frontal pole since it had larger sample size (N=250) than the other three brain regions (N ≤ 226). When performing deconvolution, we also used ROSMAP snRNA-seq as reference.

##### PsychENCODE

PsychENCODE scRNA data is processed scRNA-seq based gene expression data merged from three sources by the PsychENCODE Consortium ^21,43–45^. This data includes 15,086 genes and 4,249 cells from 35 different cell types, including microglia, astrocytes, excitatory neurons, and oligodendrocytes etc, generated using both fetal and adult brain samples. In our simulation, we only used the scRNA-seq data from adults to serve as an alternative reference panel in EPIC-unmix simulations.

#### Human peripheral blood mononuclear cells datasets (AIDA and OneK1K)

##### AIDA ^24^

The Asian Immune Diversity Atlas cohort consists of scRNA-seq data of 36266 genes from ∼1 million peripheral blood mononuclear cells (PMBCs) collected from 503 healthy Asian individuals across 22 cell types.

##### OneK1K ^5^

The OneK1K cohort consists of scRNA-seq data of 36571 genes from 1.27 million PMBCs collected from 982 healthy individuals of Northern European ancestry. Cells were classified into 14 different immune cell types across the myeloid and lymphoid lineages using a framework based on ScPred ^46^.

#### Mouse primary motor cortex cells datasets (Yao and Tasic)

##### Yao ^47^

The Yao dataset was generated by the Allen Institute including transcriptomes and epigenomes of mouse primary motor cortex cells from 538 mice. We incorporated the processed snRNA-seq dataset, generated by the 10x v2 platform then passed through QC, clustering and cell type annotation with marker genes, which contains expression profiles of 34,496 genes from 73,973 cells classified into 43 cell types.

##### Tasic ^23^

The Tasic dataset includes scRNA-seq of mouse neocortex cells isolated from VISp and ALM for 352 healthy mice of both sexes. A total of 24,411 cells were clustered into 133 cell subtypes (56 glutamatergic, 61 GABAergic and 16 non-neuronal types), and the expression profile of 34,496 genes for each cell were included.

### Other deconvolution methods

#### TCA ^12^

TCA is a frequentist deconvolution method originally designed for DNA methylation data, but also applicable for gene expression data. We applied the *TCA* R package to perform deconvolution with only bulk RNA-seq data and cell type fractions as input. Note that TCA does not need single cell reference. Genes with across-sample variance less than 10e^-8 were filtered out due to low variability.

#### bMIND ^13^

bMIND is a Bayesian deconvolution method. It first calculates a prior from scRNA-seq data, and then infers CTS expression incorporating both bulk tissue expression and the prior information via MCMC. We used the *MIND* R package with default parameters. For each dataset, the training set was chosen as the single cell prior to deconvolute the pseudo-bulk from the testing set.

### Real data based simulations

Out of the 48 individuals from the ROS/MAP snRNA-seq data, we excluded one individual with incomplete snRNA-seq results and meta data, and randomly selected 16 samples to construct pseudo-bulk, which served as target and for which we know the truth, i.e., the CTS expression profiles. The remaining 31 samples with snRNA-seq data available were used as reference to deconvolute the pseudo-bulk data. In order to assess the performance with varying reference panels, we additionally used the psychENCODE scRNA-seq ^21^ data as an alternative reference panel to deconvolute the same target pseudo-bulk data.

Besides the ROAMAP based simulations, we also used sc/snRNA-seq data from two more tissues to further evaluate the performance of EPIC-unmix. We paired the Yao and Tasic datasets from mouse brain tissue, subsetted and regrouped glutamatergic cells of 5 subtypes/layers: L2/3 IT, L4/5 IT, L5 PT, L6 CT, and L6 IT, for each of the two datasets separately. We only retained genes that were expressed in at least 80% of the target pseudo bulks generated by Yao. The final datasets contained 13,947 genes and 36,920 (Yao) vs. 10,480 (Tasic) cells of the 5 subtypes.

For the two human blood data, we down-sampled the OneK1K dataset, retaining cells from 10% randomly selected donors (n = 90) for computational efficiency. We paired it with the AIDA dataset and selected natural killer cells, memory B cells, naive B cells, gamma-delta T cells, and regulatory T cells, to generate pseudo bulk and perform deconvolution. After grouping AIDA cells by donors as the target pseudo bulks, we similarly only retained genes that were expressed in at least 80% of the samples. The final datasets contained 10,283 genes and 49,841(AIDA) vs.120,765 (OneK1K) cells of the 5 cell types.

For the datasets selected for pseudo bulk generation (AIDA and Yao), we randomly partitioned the cells, grouped by donors, into a training set and a testing set. Cells from the testing set were aggregated as pseudo bulks by their donor ID. For cells within each pseudo bulk, we used their scRNA-seq profiles and cell type information to calculate sample-level bulk expression per gene, cell type proportions, and the true CTS expression for each gene in each cell type. Given the sparsity of snRNA-seq data, an additional step of QC was performed during the generation of pseudo bulks, where we only retained genes that were expressed in more than 80% of the pseudo bulks.

We applied EPIC-unmix and two alternative methods (TCA and bMIND) on the simulated bulk RNA-seq samples for CTS deconvolution. All methods deconvolute the two-dimensional (genes by samples) bulk RNA-seq data as the mixture of various cell types into a three-dimensional tensor of CTS gene expressions (genes by samples by cell types). As each method retains predictions of different genes through distinct gene filtering strategies, we focused on the intersection of their retained genes, and used the absolute value of Pearson Correlation Coefficient (PCC) of the true and predicted CTS expression of those genes to evaluate the performance of different methods.

### Gene selection strategy

As it is impossible to accurately estimate expression levels for every gene in every cell type ^11,13^, we developed gene selection strategies to prioritize genes that are more likely to be well-inferred in each cell type. To obtain such lists of high-confidence genes, we developed two types of gene selection strategies based on different purposes of simulation studies and downstream analysis. These strategies showed promising results in simulations.

In simulations using human blood and mouse brain datasets, we obtained the list of highly expressed genes that were expressed in >80% of the pseudo bulks in the “training set”. We then applied “FindMarkers” in Seurat v4.1.0 on the “testing set”. We selected genes with Seurat differential expression p-value < 0.05 and top 200 (at maximum) genes based on log2(fold change, FC) for each cell type. We then intersected the genes with those passing bulk-level QC as the prioritized gene sets for prediction and model performance evaluation (**Figure S4A**).

In simulations using the ROSMAP human brain data, selected genes were constructed by taking the union of two sets of genes **(Figure S4B**). The first set included genes showing reasonable agreement among three independent datasets, including CTS bulk expression ^20^ and two snRNA-seq ^16,17^ datasets. For each cell type in each pair of datasets, we first fitted a simple linear regression model and then retained only genes where the differences between observed and predicted in one dataset is within 0.5*observed value of the other dataset. This set was then reduced by keeping only genes with normalized snRNA-seq raw counts > 0.5 in the ROSMAP snRNA-seq dataset ^6^ in each cell type. We further normalized the expression levels by scaling the size factors in scran ^48^. Finally, we further reduced this set by only retaining cell type marker genes with Seurat differential expression p-value < 0.05 and top 200 (at maximum) genes based on log2(FC), using “FindMarkers” in Seurat v4.1.0 following the standard pipeline with default parameters ^49^ in each cell type. All analyses were performed in R/4.1.0. The second set included marker genes reported in literature ^18,19^.

### CTS differentially expressed gene (DEG) analysis

After obtaining EPIC-unmix inferred sample-level CTS expression profiles for ROSMAP or MSBB samples, we conducted DEG analysis separately for each gene in each cell type by fitting a logistic regression model of binary AD status (1=case, 0=control) against gene expression, adjusting for covariates including sex, age of death, pmi, education year for ROSMAP and sex, race, age of death and pmi for MSBB.

We then meta-analyzed the DEG results from the two datasets using METAL^25^. For each cell type, the list of DEGs revealed from meta-analysis (FDR < 5%) was subject to Gene Ontology enrichment analysis, with R package clusterprofiler ^50^.

### CTS cis-eQTL analysis

Post EPIC-unmix inference, we also performed eQTL analysis using WGS data, considering SNPs with MAF > 0.01 in the +/- 1MB region of each gene body. We used MatrixeQTL ^51^ for cis-eQTL analysis with a linear model, correcting for covariates, sex, age of death, pmi, education year, final consensus cognitive diagnosis (COGDX) for ROSMAP and sex, race, age of death, pmi, clinical dementia rating scale (CDR) for MSBB.

We assessed our CTS eQTL results using two approaches. First, we compared our statistically significant CTS eQTL-gene pairs (FDR < 0.05) with those from external human brain CTS eQTL datasets. We examined the number of shared/exclusively identified pairs between our results and Bryois et al ^29^, and also checked directions and magnitudes of eQTL effect sizes for the shared gene-variant pairs. For those identified exclusively by one, we additionally leveraged two microglia eQTL datasets ^30,31^ to examine the overlaps. Beyond comparing to existing CTS eQTL studies, we also overlapped our microglia-specific cis-eQTL variants with microglia ATAC-seq and pcHi-C loops ^20^, and AD GWAS variants ^36^ to highlight potential biologically interesting variant/genes. We presented visualizations in the WashU genome browser ^52^. All the analyses were performed in R/4.1.0.

### Meta-analysis

To increase the sample size, we conducted meta-analysis using METAL ^25^ to combine CTS DEG and cis-eQTL derived from ROSMAP and MSBB. For DEGs, we used inverse variance weights to calculate meta z-scores and p-value based on effect sizes and standard errors of each gene from the two studies. For cis-eQTLs, we used sample size weighted scheme to derive meta z-scores and p-values based on directions of effect size and original p-values of the two studies. To account for multiple testing, we used Benjamini & Hochberg procedure implemented by “p.adjust” in R to calculate the false discovery rate (FDR), with 5% cutoff to define significance.

## Acknowledgement

This study was supported by NIH grant R01AG079291, U01DA052713 and RF1AG082938. We thank the ROSMAP and MSBB participants. QS was supported by a TOPMed Fellowship. JW was partially supported by T32ES007018. YL was partially supported by P50HD103573 and U24AR076730.

