## Supplementary material for "Cell-type specific inference from bulk RNA-sequencing data by integrating single cell reference profiles via EPIC-unmix": Figure S1

### Supplementary Figures

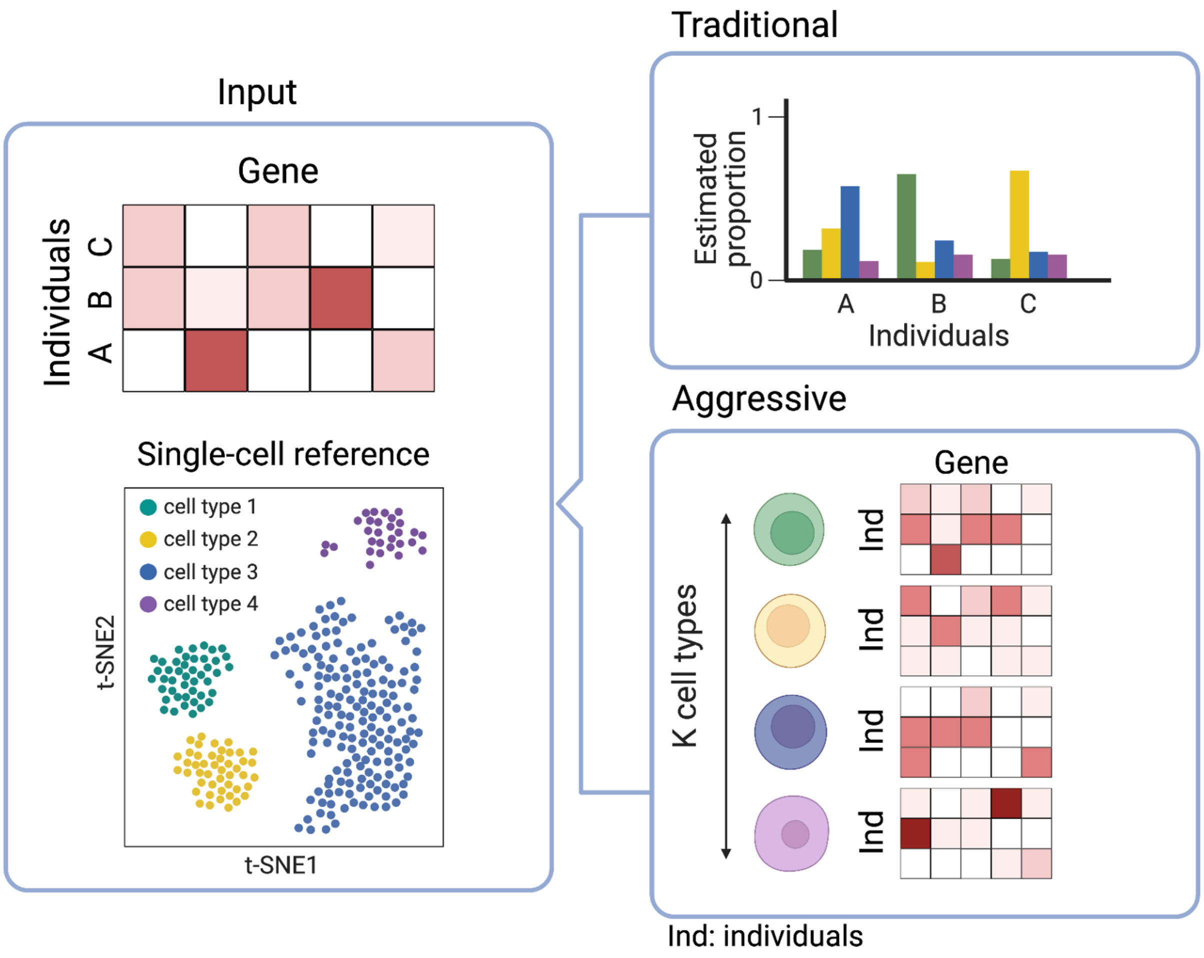


**Figure S1. Cell type deconvolution methods overview and classifications.** For illustration brevity, we consider three individuals A, B, C, and five genes. Top left panel illustrates input bulk RNA-seq data for study samples while bottom left shows single cell reference data. With the two pieces of input on the left, we consider methods as "traditional" if they provide only cell-type fractions for each study sample (top right panel). In contrast, methods that generate sample-level CTS profiles are classified as "aggressive" (bottom right panel).

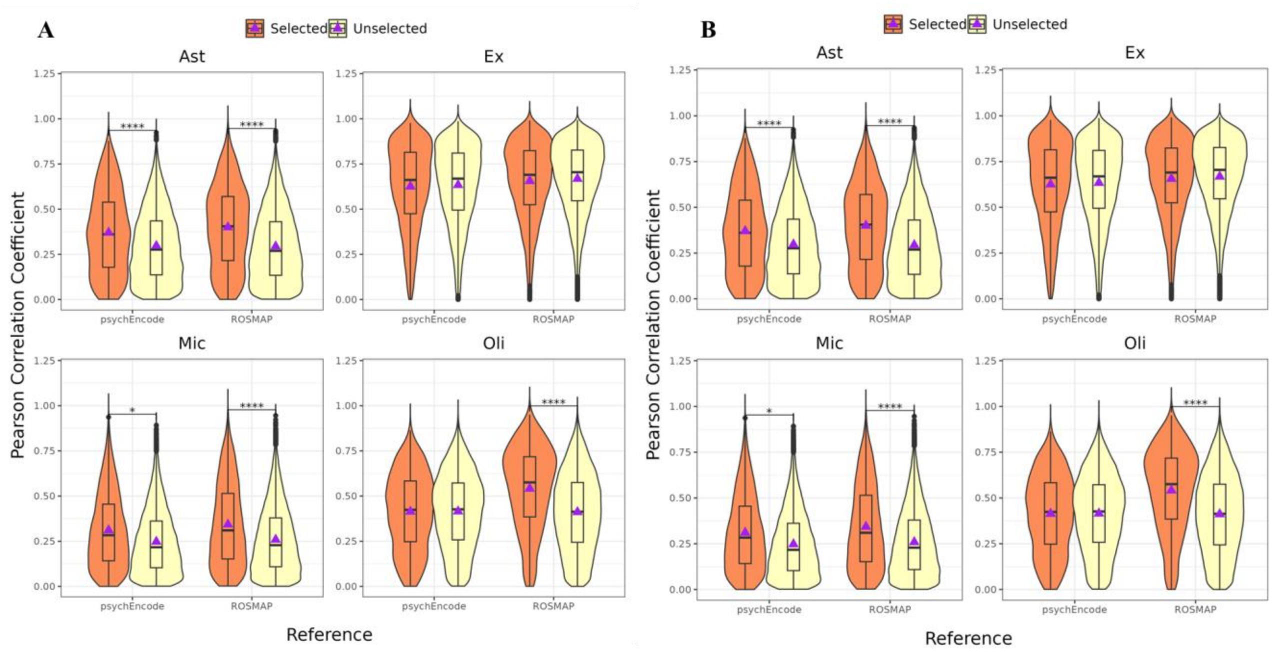


**Figure S2. Deconvolution performance comparison between selected and unselected genes using bMIND and TCA.** Performance, measured by the absolute Pearson correlation between inferred CTS expression and truth, is compared for selected and unselected genes in ROSMAP-based simulations using bMIND (**A**) and TCA (**B**) for deconvolution.

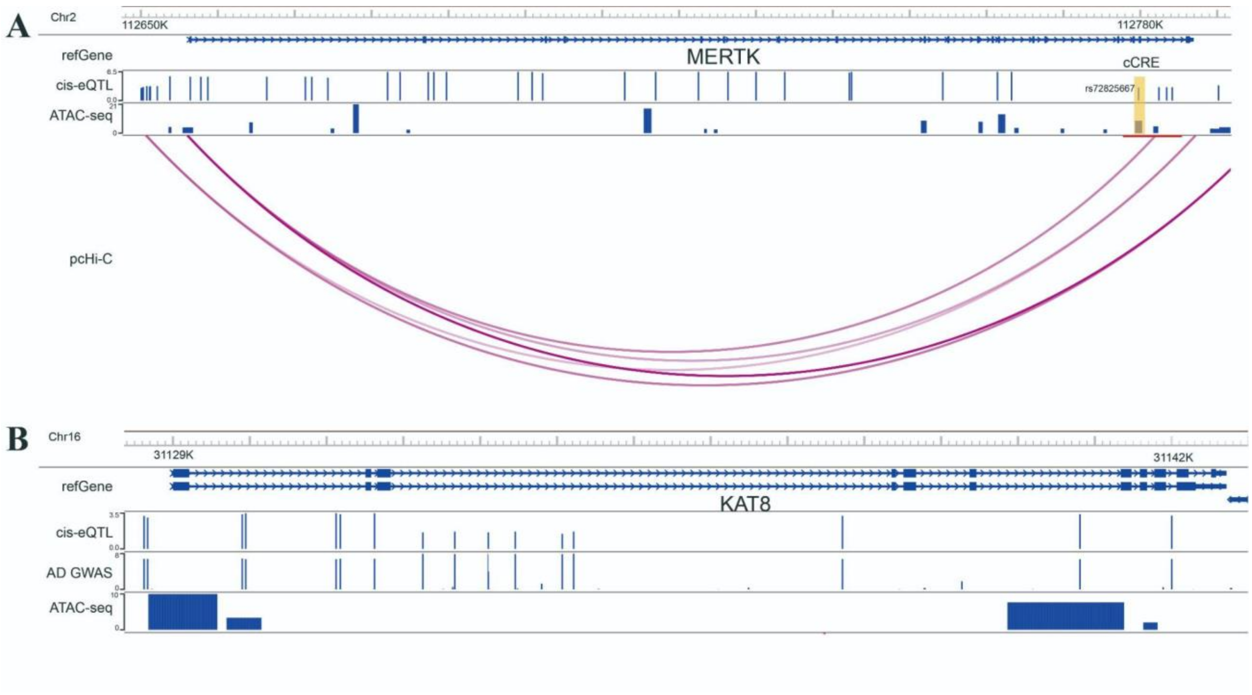


**Figure S3. Highlights of microglia specific eQTLs for *MERTK* and *KAT8*. A.** Annotated Genome browser view of *MERTK*. Tracks from top to bottom are: genome position with GRch37 build; gene location; *MERTK's* CTS cis-eQTL (height of signal = $-\log_{10}$(adjusted p-value)); microglia specific ATAC-seq narrow peaks (height of signal = score); microglia specific pcHiC loops. **B.** Annotated Genome browser view of *KAT8*. Tracks from top to bottom are: genome position with GRch37 build; gene location; *KAT8's* CTS cis-eQTL (height of signal = $-\log_{10}$(adjusted p-value)); AD GWAS variants [1] (height of signal = $-\log_{10}$ (GWAS p-value)); microglia specific ATAC-seq narrow peaks (height of signal = score).

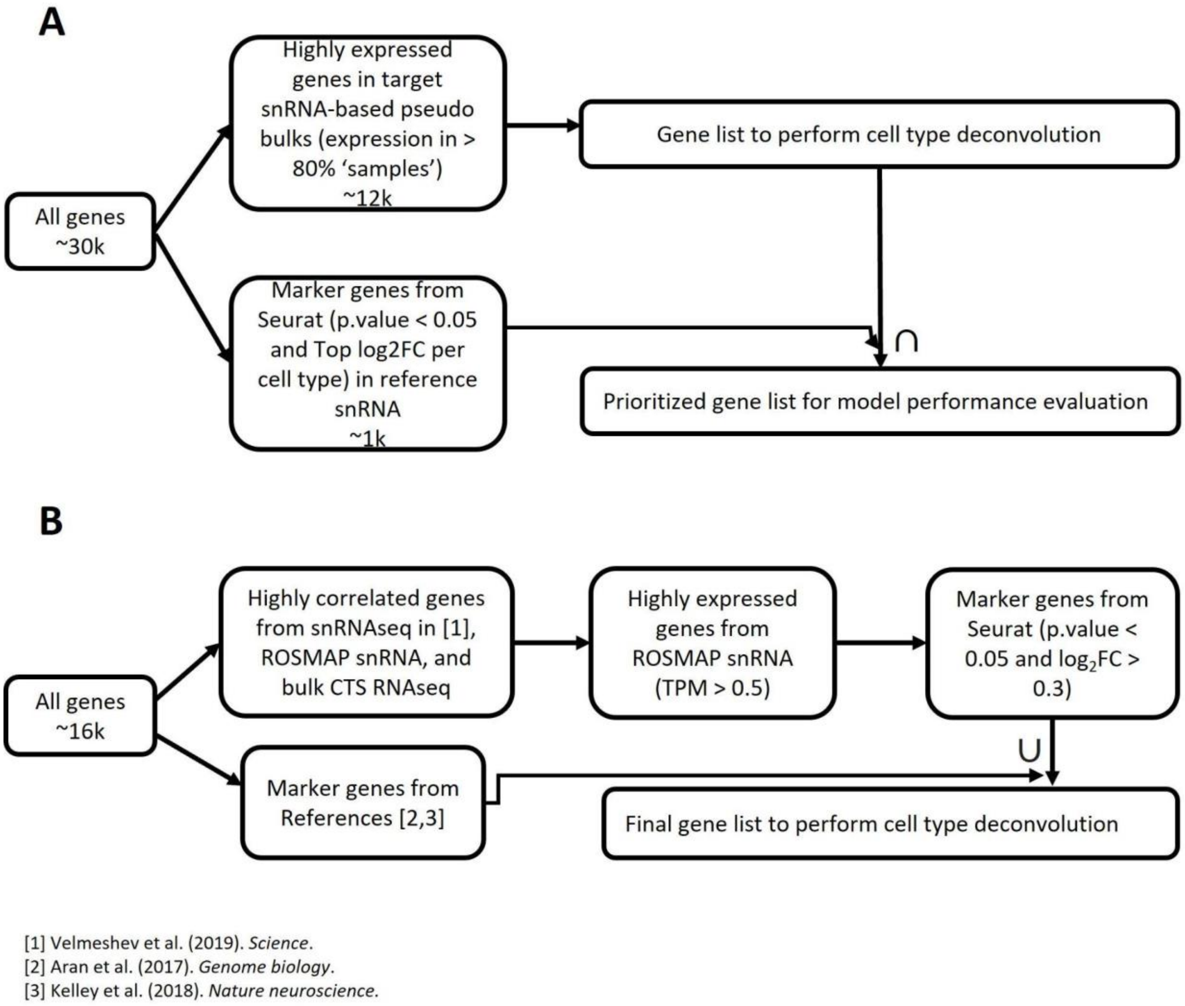


**Figure S4. Illustration of gene selection strategies. A.** Gene selection strategy for human blood and mouse brain snRNA-seq based simulations study of deconvolutional methods. **B.** Gene selection strategy for ROSMAP human brain snRNA-seq based simulations study of deconvolutional methods.
